# Rapid *in situ* mutation detection in extracellular vesicle-DNA

**DOI:** 10.1101/2024.02.26.582068

**Authors:** Md. Mofizur Rahman, Lixue Wang, Md. Motiar Rahman, Yundi Chen, Wenlong Zhang, Jing Wang, Luke P. Lee, Yuan Wan

## Abstract

A PCR- and sequencing-free mutation detection assay facilitates cancer diagnosis and reduces over-reliance on specialized equipment. This benefit was highlighted during the pandemic when high demand for viral nucleic acid testing often sidelined mutation analysis. This shift led to substantial challenges for patients on targeted therapy in tracking mutations. Here, we report a 30-minute DNA mutation detection technique using Cas12a-loaded liposomes in a microplate reader, a fundamental laboratory tool. CRISPR-Cas12a complex and fluorescence-quenching (FQ) probes are introduced into tumor-derived extracellular vesicles (EV) through membrane fusion. When CRISPR-RNA hybridizes with the DNA target, activated Cas12a can *trans*-cleave FQ probes, resulting in fluorescence signals for the quantification of DNA mutation. Future advancements in multiplex and high-throughput mutation detection using this assay will streamline self-diagnosis and treatment monitoring at home.

## 1 Introduction

Growing evidence has shown the clinical relevance of extracellular vesicles (EV) in cancer. Recent studies have validated the use of EV analysis for (early) cancer diagnosis, treatment monitoring, and prognosis [1, 2]. In the realm of EV-based cancer diagnostics, two primary technical approaches are currently used [3]. One approach involves on-chip analysis, where EVs are enriched on surfaces for the profiling of proteins in EV membranes. This method is convenient but merely analyzes membrane proteins. In contrast, the analysis of encapsulated EV cargos can be advantageous for EV-based cancer diagnosis [4]. However, the protective lipid bilayer of EVs impedes the access of detecting reagents to the cargo, rendering cargo detection challenging through the existing on-chip analysis. Alternatively, the second approach focuses on the isolation of EVs. Subsequently, multiple processing steps are required to properly extract and purify cargo molecules for molecular analysis. This approach enables comprehensive and accurate analysis. However, the labor-intensive sample preparation procedure significantly impedes analysis efficiency and promptness. Ideally, an on-chip detection approach that can circumvent cargo extraction and purification but interrogate wrapped cargo is highly desired.

On the other hand, cancer patient services encountered significant challenges amidst the COVID-19 pandemic. Visits to cancer centers were minimized to reduce exposure risks, yet these protective measures also constrained cancer diagnosis and surveillance. This dilemma was apparent in patients undergoing mutation-targeted therapy, who could benefit from resistance tracking and timely medication adjustment [5-7]. In addition, genetic testing for cancers was crowded by the overwhelming COVID-19 nucleic acid tests, as both primarily rely on PCR and next-generation sequencing (NGS) [8, 9]. In brief, the pandemic severely impacted cancer diagnosis and treatment monitoring [6, 10, 11]. This challenge highlights the critical importance of promptness and convenience in genetic testing. Here, we report a PCR-, NGS-, and *in situ* amplification-free liquid biopsy using tumor-derived extracellular vesicles (EV) for DNA mutation detection in 30 minutes [7, 12-14]. Tumor-derived EVs in plasma are immunocaptured onto surfaces followed by membrane fusion with liposomes (LP) containing CRISPR-Cas12a and fluorescence-quenching (FQ) probes (Supplementary Table 1). The hybridization between DNA targets and complementary CRISPR RNAs (crRNA) triggers the cleavage activity of Cas12a, enabling the detection of DNA mutation through fluorescence signals emitted from shredded FQ probes (Fig. 1).

**Figure 1:**
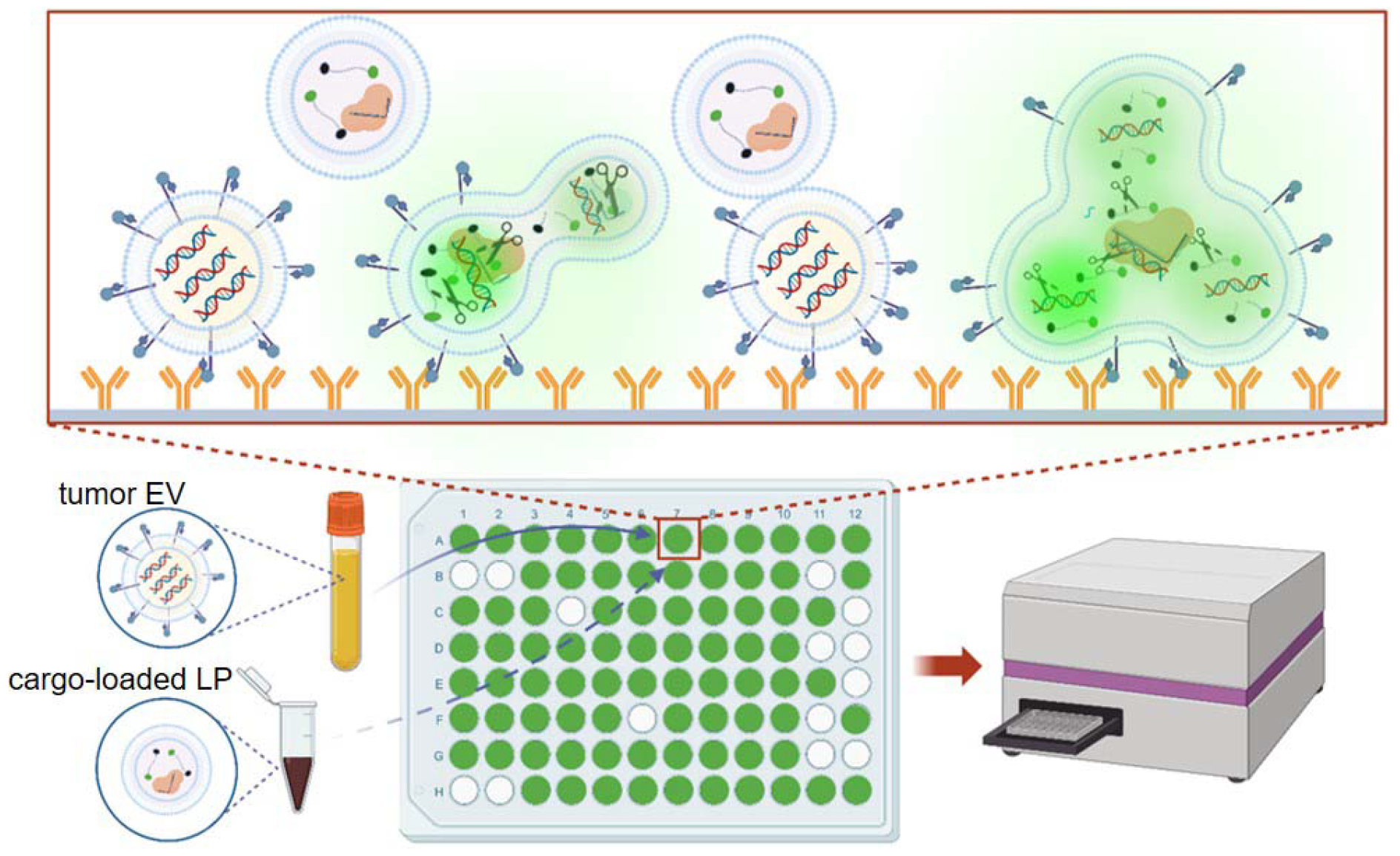
Schematic of CRISPR-Cas12a-based EV-DNA mutation detection. EVs in plasma are immunocaptured onto anti-EpCAM antibody-grafted surfaces followed by membrane fusion with LPs containing Cas12a-crRNA and FQ probes. When crRNA hybridizes with the DNA target, activated Cas12a can cleave FQ probes, resulting in fluorescence signals. Therefore, DNA mutation can be qualitatively detected with a typical microplate reader.

## 2 Materials and methods

### 2.1 Preparation of liposomes

Liposomes were prepared using DOTAP, DMPC, DOPE, and Cholesterol in molar ratios of 550:100:200:100, with 1 mol% NBD-PE and 1 mol% Rhodamine-PE. The thin film hydration method was employed to create a lipid layer, evaporating the organic solvent at 50°C with a Rotavapor (30 rpm). Argon was used to further remove solvent in a fume hood for one hour. The lipid film in the tube was vacuum-dried overnight. To make liposomes loaded with CRISPR-Cas12a complex and FQ probe, the lipid film was hydrated at room temperature, and the cargo was added to the tube. Control groups were prepared similarly. The mixture was briefly vortexed for 5 sec, stirred at 100 rpm for 2 h at room temperature, and then left overnight at 4°C. Finally, 100 nm liposomes were obtained by extruding the liposomes 20 times through a 0.1-micron polycarbonate membrane using an Avanti Mini-Extruder, and any free reagents were removed with a 300-KDa membrane filter following the manufacturer’s instructions.

### 2.2 Cell culture and isolation of extracellular vesicles

H441 and H1975 were purchased from ATCC. Cells passed the mycoplasma contamination test, and were cultured in DMEM (Corning, USA). All mediums were supplemented with 5% (v/v) EV-depleted Fetal Bovine Serum (FBS) (Thermal Fisher, USA), 100 units/ml penicillin, /ml streptomycin, and 1% non-essential amino acid. All cell lines were incubated at 37 °C with 5% CO_2_ and a 95% humidified atmosphere. Cells at 70% confluency were cultured in FBS-free medium for 48 h. The supernatant was then centrifuged at 2,500 g at 4°C for 15 min, followed by a second centrifugation at 16,500 g for 20 min. Subsequently, the medium was filtered through a 0.22-μm pore filter, and the supernatant was ultracentrifuged at 100,000 g at 4°C for 4 h.

### 2.3 Nanosight measurement

Liposomes and EV samples were suspended in 200 μl of PBS. Size distribution of liposome and EV were measured with Nanosight NS300 according to manufacturer’s instructions.

### 2.4 Zeta potential measurement

Zeta potential measurement was conducted using a Zetasizer Nano ZS system. Approximately 10 μl of the sample in 990 μl of DI water was transferred to a Malvern Clear Zeta Potential cell. Three independent aliquots were analyzed, and three measurements were taken for each aliquot.

### 2.5 Transmission electron microscope

For TEM, 5 μl of the sample was applied to a Formvar-coated copper grid (400-mesh) and incubated for 3 min at room temperature. Excess sample was removed with filter paper, and then the grid was negatively stained with 1% aqueous uranyl acetate for 1 min. After blotting dry, the samples were examined using a FEI Tecnai transmission electron microscope at 100 kV.

### 2.6 FRET assay

FRET analysis was conducted using a TECAN Spark. For the NBD-chol/Rhod-PE pair, the excitation wavelength was set at 460 nm, and emission spectra were recorded from 500 to 700 nm. In a 96-well black-walled microplate, liposomes labeled with both NBD-PE and Rhod-PE were mixed with unlabeled EVs at liposome to EV ratios of 1:1, 1:3, and 1:5, totaling 50 μl. The EV-liposome fusion reaction mixture was incubated up to 15 min, and fusion efficiency was determined by measuring the change in NBD fluorescence intensity before and after fusion. After fusion, 10% Triton X-100 (Sigma Aldrich, X100) was added to dissolve all vesicles and obtain the NBD fluorescence intensity *F* _∞_ . The fusion efficiency was calculated using the formula: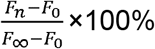, where *F*_0_ and *F*_*n*_ represent the fluorescence intensities before and after fusion, respectively.

### 2.7 In Vitro Cas12a-based DNA mutation detection

The reaction mixture consisted of 50 nM Cas-12a, 50 nM crRNA, 10 U RNase inhibitor, 40 nM FQ probe. This mixture was combined with the specified concentration of reaction buffer (50 μL containing 40 mM Tris-HCl at pH 7.5, 60 mM NaCl, and 6 mM MgCl_2_) and then incubated at 37°C for 30 min. Fluorescent intensities were measured using a Tecan multi-plate reader.

### 2.8 Fluorescence microscope

Isolated EVs were stained with PKH26 at 37°C for 30 min. Excess dye was removed using an ultracel-30 membrane (Millipore, MRCF0R030). The labeled EVs were then collected and resuspended in PBS. Fluorescence images were captured using an Olympus IX83 microscope.

### 2.9 Mutation detection in a microplate

In the Cas12a assay, a black-walled 96-well ELISA plate coated with anti-EpCAM antibodies was incubated at room temperature with 100 μl of purified EVs or plasma for EV capture. After three washes with PBS containing 0.05% Tween 20, sample wells were incubated with 50 μl of a reaction solution containing cargo-loaded liposomes. The fluorescent signal was then measured using a microplate reader. A positive EV assay result was defined as a signal equal to or greater than a cut-off threshold, which was determined as the mean signal of the negative control samples plus three times their standard deviation.

### 2.10 Collection of plasma samples

Blood samples were obtained at the Second Hospital of Nanjing according to an institutional-review-board-approved protocol (IRB: 2016-LY-KT038). Samples were drawn into 10-ml EDTA (K2) tubes (Vacutainer; Becton Dickinson) from peripheral venepuncture. After centrifugation at 2,500 g for 15 min, plasma was collected and filtered using a 0.22-μm filter.

### 2.11 Statistical analysis

Quantitative results were presented as mean±SD. Student’s unpaired t-test was used to compare control treated samples against experimental samples. For multiple treated groups, statistical significance was examined using one-way ANOVA.

## 3 Results and discussion

Fusogenic LPs [15], tailored for delivering Cas12a-crRNA complex and FQ probes [16], were formulated with an average size of 100 nm, aligning with the size of small EVs (Fig. 2a). The average zeta potential of LPs, Cas12a-crRNA complex-loaded LPs, H1975 EVs, and H441 EVs were 71.02 mV, 40.73 mV, -46.31 mV, ad -44.25 mV, respectively. These LPs engaged in membrane fusion with EVs through electrostatic interactions, resulting in fused vesicles with sizes ranging from 200 nm to 800 nm. The complete fusion and fusion intermediate were visualized by electron microscopy images (Fig. 2b and Supplementary Fig.1). After fusion, the average zeta potential of H1975 EV-LP and H441 EV-LP was -10.65 and -11.33 mV, respectively. Förster resonant energy transfer (FRET) assay also verified their fusion [17, 18]. Modulating the EV-to-LP ratio, either by increasing or decreasing it, reduced FRET efficiency (Fig. 2c), suggesting that membrane fusion scattered FRET dyes on liposomal membranes. Nevertheless, the average fusion efficiency was determined to fall within the range of 50% to 90%.

**Figure 2:**
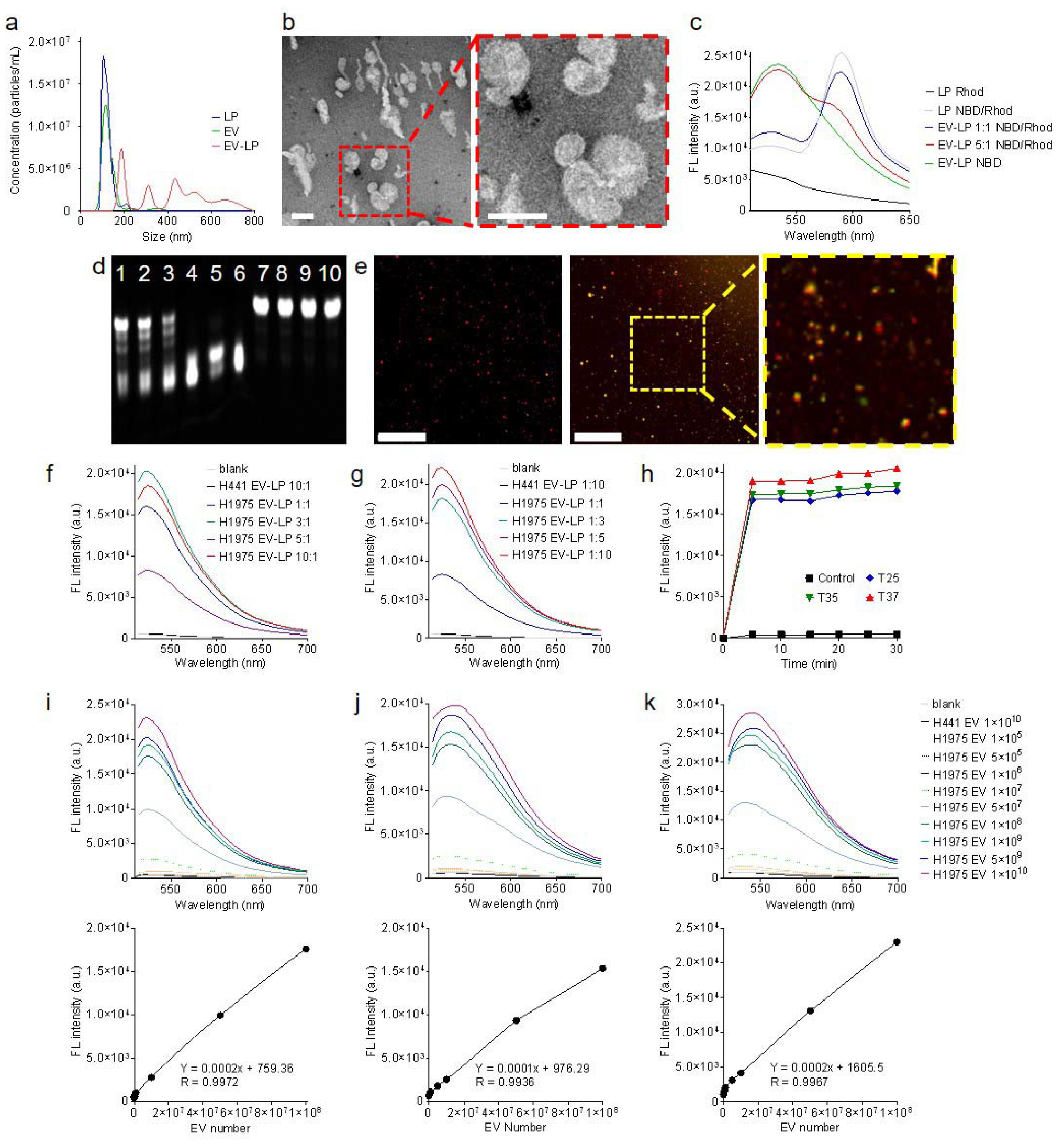
Characterization and optimization of Cas12a-based assay. **a**, Size distributions of LPs, EVs, and fused EV-LP. **b**, Electron microscope image of an EV-LP fusion (scale bar: 100 nm). **c**, Fluorescence signals of dyes-labeled LPs in the FRET assay. **d**, Gel analysis of the FQ probes after cleavage (Lane 1-5: FQ probes were *trans*-cleaved by Cas12a in the presence of L858R mutation; H1975 EV amount was 1×10^5^, 1×10^6^, 1×10^7^, 1×10^8^, and 1×10^9^, respectively. Lane 6: FQ probes were cleaved by Cas12a, which was used as a positive control. Lane 7: FQ probes were not cleaved by Cas12a in the absence of L858R mutation; H441 EV amount was 1×10^10^. Lane 8: FQ probes were not cleaved by Cas12a in the absence of EV DNA. Lane 9: FQ probes were not cleaved in the absence of Cas12a. Lane 10: FQ probes only. **e**, Fluorescence co-localization analysis of fused EV-LP. In the absence of L858R mutation, only fluorescence signals emitted from PKH26 (red) were observed in H441 EV-LP (left; scale bar: 50 μm). In the presence of L858R mutation, dual fluorescence signals emitted from PKH26 and shredded FQ probes (green) were observed in H1975 EV-LP (right). The insert shows a local zoom-in view. **f-g**, Quantitative fluorescence intensity detected from cleaved FQ probes in various EV-to-LP ratios. The reaction volume was 50 μl. **h**, Quantitative fluorescence intensity detected from EV-to-LP ratio of 5:1 after 30-min incubation at 25, 35, and 37 □°C, respectively. The reaction volume was 50 μl. **h**, LOD of L858R mutation in H1975 EVs suspended in PBS. **i**, LOD of L858R mutation in H1975 EVs suspended in PBS followed by immunocapture onto surfaces. **j**, LOD of L858R mutation in H1975 EVs suspended in human plasma followed by immunocapture onto surfaces.

The assay was optimized with tumor EV model samples. Mutation detection specificity was confirmed before performing *in situ* detection. Gel electrophoresis verified concentration-dependent *trans*-cleavage of FQ probes into short fragments upon exposure to activated Cas12a in the presence of the L858R mutation. In contrast, this cleavage was absent in scenarios devoid of this mutation (Fig. 2d). In parallel, we determined that the average loading efficiency of Cas12a-crRNA and FQ probe was 89.22% and 94.67%, respectively. Next, fluorescence co-localization analysis demonstrated the delivery of cargo by LPs to PKH26-labeled EVs. A substantial green fluorescence signals emitted from cleaved FQ probes were observed in ∼85% of H1975 EVs harboring the L858R mutation. In contrast, less than 0.1% of H441 EVs lacking the L858R mutation exhibited negligible green fluorescence signals (Fig. 2e). The quantitative analysis using a microplate reader revealed that the signal intensity exhibited a proportional increase with the rising level of L858R copies, eventually reaching saturation (Fig 2f and Supplementary Fig. 2). In this scenario, all Cas12a-crRNA complexes effectively engaged and bound to the targeted DNA. The Cas12a *trans*-cleaved a limited number of FQ probes. A parallel trend was observed when maintaining the L858R copies but increasing the quantity of Cas12a-crRNA complexes (Fig 2g). We postulated that all mutant targets underwent *cis*-cleavage, while an excess of FQ probes in fused vesicles simultaneously experienced *trans*-cleavage. The findings indicated that the overall quantity of FQ probes in fused vesicles determined the signal intensity. Moreover, we observed that the reaction kinetics of Cas12a remained relatively consistent at 25 °C compared to those at 35 °C and 37 °C. Maximum signal intensities were detected after a mere 5-minute incubation (Fig 2h). To facilitate this study, we adopted 25 °C in the subsequent experiments. Notably, if conditions permit, 37 °C is indeed the optimal choice for mutation detection.

The fluorescence threshold (FLt) and limit of detection (LOD) of this assay were measured. Each H1975 EV contains ∼10.5 to 24.6 copies of DNA fragments harboring L858R mutation. Various amounts of H1975 EVs were resuspended in PBS and subjected to detection using 2.5×10^8^ LPs. The FLt was established at 538 arbitrary units (a.u.), and the LOD^PBS^ in solution was calculated to be 3.14×10^5^ H1975 EVs. Using the same experiment setup, we found that FLt and LOD^PBS^ on the EpCAM-coated plate surface were 642 a.u. and 6.27×10^5^ H1975 EVs. In addition, we tested 5 μl of EV-spiked human plasma samples to simulate clinical conditions. The average immunocapture efficiency was 92.45% in 15 minutes. The FLt and LOD^Plasma^ on the anti-EpCAM antibody-coated plate surface was 1,065 a.u. and 8.3×10^5^ H1975 EVs (Fig. 2i-k and supplementary Fig. 3). Notably, mutation detection was fulfilled in 30 minutes. In comparison, real-time quantitative PCR (RT-qPCR) spent approximately two hours detecting mutation from ∼8×10^5^ H1975 EVs (Supplementary Fig. 4). These findings demonstrate the feasibility, accuracy, and reliability of this assay. Lastly, we assessed this assay using plasma samples collected from 10 healthy donors and 30 patients with late-stage non-small cell lung cancer (Supplementary Fig. 5 and Supplementary Table 2). Cancer patients’ L858R mutation in circulating tumor DNA was verified with RT-qPCR. We determined that the sensitivity, specificity, and accuracy of our assay was 86.7%, 90%, and 87.5%, respectively.

## 4 Conclusion

RT-qPCR and NGS are standard diagnostic tools in clinical laboratories [19-21]. Both require specialized equipment and skilled staff [22]. To monitor mutation status of patients undergoing EGFR-targeted therapy, RT-qPCR takes a minimum of two hours, and NGS takes at least two days (Supplementary Table 3). In contrast, this CRISPR-Cas12 assay does not demand specialized sample handling or complex data analysis [23]. It achieves mutation detection using a common microplate reader in 30 minutes, reducing the duration and exposure risks of vulnerable patients with advanced cancers in hospitals. The detection sensitivity and specificity are on par with those of RT-qPCR. Notably, this assay eliminates the need for *in situ* amplification of nucleic acids [24], streamlining the entire detection process for increased simplicity and efficiency. In future, by using a cocktail of Cas12a-crRNAs targeting multiple mutations, we can conduct simultaneous and high-throughput detection of various actionable mutations for EGFR-targeted therapy. Moreover, the design of this assay has the potential for development into cost-effective point-of-care devices for bedside diagnosis in the ward or even patient self-diagnosis from home, facilitating the monitoring of treatment effectiveness out of the oncology centers. In summary, our assay offers a competitive alternative to RT-qPCR and NGS for mutation detection, boosting self-diagnosis and treatment monitoring.

## Abbreviations

FQ: Fluorescence-quenching
EV: Extracellular vesicles
NGS: Next-generation sequencing
LP: Liposome
FBS: Fetal Bovine Serum
FRET: Förster resonant energy transfer
FLt: Fluorescence threshold
LOD: Limit of detection
RT-qPCR: Real-time quantitative PCR

## Acknowledgements

No applicable

## Autho contributions

Y.W., J.W., and L.L. designed the project. Mofizur R., L.W., Motiar R., Y.C, and W.Z. performed the experiments, collected, and analyzed the data. All authors contributed to the writing of the manuscript, discussed the results and implications, and edited the paper at all stages.

## Availability of data and materials

In supporting information, sequences of oligomers, patients’ information, comparison Cas12a-based assay with PCR and NGS, TEM images, fluorescence co-localization analysis images, fluorescence threshold of mutation detection, PCR detection of mutation, and DNA mutation detected in patient’s samples were provided.

## Declarations

### Ethics approval and consent to participate

Blood samples were obtained at the Second Hospital of Nanjing according to an institutional-review-board-approved protocol (IRB: 2016-LY-KT038).

