## Supplementary material for "Rapid *in situ* mutation detection in extracellular vesicle-DNA": SI

**Supplemental Table S1. Sequences of oligomers used in this study.**

| Name | Sequence (5’ to 3’) |
| --- | --- |
| crRNA for *EGFR* L858R | /AlTR1/rUrArArUrUrUrCrUrArCrUrArArGrUrGrUrArGrArUrGrCrCrCrGrCrCrCrArArArArUrCrUrGrUrGrArUrCrU/AlTR2/ |
| FQ probe | FAM-TTTTAATTTT-IABkFQ |

**Supplemental Table S2. Clinical information of patients with stage-IV NSCLC (TP: true positive, FP: false positive, TN: true negative, and FN: false negative).**

| Well | RT-qPCR | Cas12a-based detection | Sex | Age |
| --- | --- | --- | --- | --- |
| A2 | L858R+ | FN | M | 68 |
| A3 | L858R+ | TP | M | 77 |
| A4 | L858R+ | TP | M | 55 |
| A5 | L858R+ | TP | F | 62 |
| A6 | L858R+ | FN | F | 58 |
| A7 | L858R+ | TP | M | 57 |
| A8 | L858R+ | TP | F | 52 |
| A9 | L858R+ | TP | F | 73 |
| A10 | L858R+ | TP | F | 64 |
| A11 | L858R+ | FN | M | 45 |
| B2 | L858R+ | TP | M | 28 |
| B3 | L858R+ | TP | F | 45 |
| B4 | L858R+ | TP | M | 75 |
| B5 | L858R+ | TP | F | 77 |
| B6 | L858R+ | TP | M | 51 |
| B7 | L858R+ | TP | M | 56 |
| B8 | L858R+ | TP | M | 58 |
| B9 | L858R+ | TP | M | 72 |
| B10 | L858R+ | TP | F | 70 |
| B11 | L858R+ | TP | F | 81 |
| C2 | L858R+ | TP | F | 70 |
| C3 | L858R+ | TP | M | 70 |
| C4 | L858R+ | TP | M | 65 |
| C5 | L858R+ | TP | F | 67 |
| C6 | L858R+ | FN | M | 73 |
| C7 | L858R+ | TP | F | 68 |
| C8 | L858R+ | TP | F | 56 |
| C9 | L858R+ | TP | F | 69 |
| C10 | L858R+ | TP | M | 48 |
| C11 | L858R+ | TP | F | 61 |
| D2 | L858R- | TN | M | 54 |
| D3 | L858R- | TN | M | 52 |
| D4 | L858R- | FP | M | 63 |
| D5 | L858R- | TN | F | 61 |
| D6 | L858R- | TN | M | 49 |
| D7 | L858R- | TN | F | 74 |
| D8 | L858R- | TN | F | 56 |
| D9 | L858R- | TN | M | 50 |
| D10 | L858R- | TN | M | 33 |
| D11 | L858R- | TN | M | 51 |

**Supplemental Table S3. Comparison Cas12a-based assay with RT-qPCR and NGS.**

|  | NGS | RT-qPCR | CRIPSR-Cas12a assay |
| --- | --- | --- | --- |
| Plasma isolation | 10 min | 10 min | 10 min |
| EV isolation | 15 min – 4 h | 15 min – 4 h | 15 min |
| DNA preparation | >9 h | ~30 min | - |
| Detection | >6 h | ~1 h | ~5 min |
| Data analysis | >8 h | >10 min | instaneous |
| Turnaround time | >2 day | >2 h | ~30 min |
| LOD of mutation | 0.1-1% | 0.1-5% | ~1% |
| Bulky instrument | Yes | Yes | Yes or No |
| Cost | High | Medium | Low |

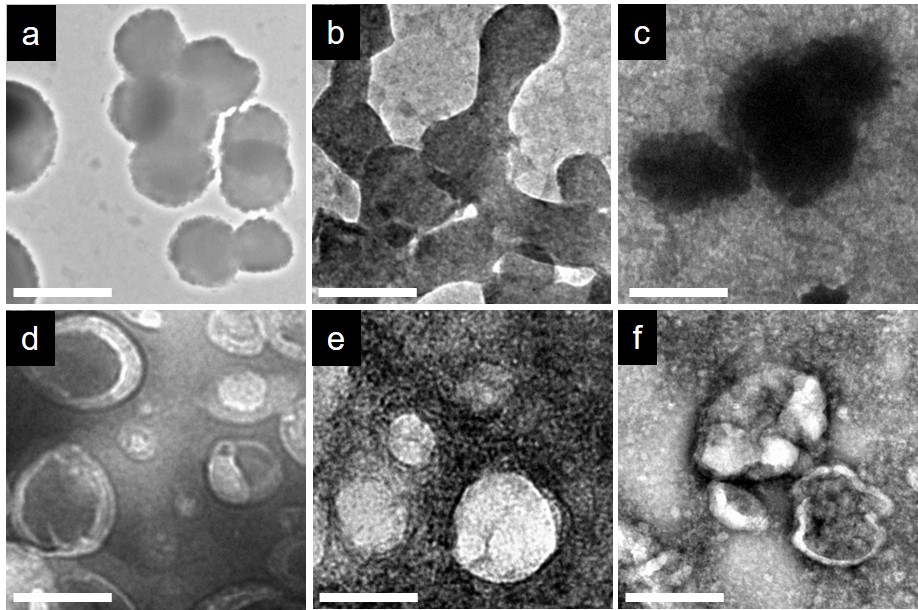

**Figure S1. Representative TEM images of EV, LPs, and EV-LPs**. **a-c**, EV-LP multiple fusion and fusion intermediate. **d**, LPs only. **e**, H1975 EVs. **f**, H441 EVs. Scale bar is 100 nm.

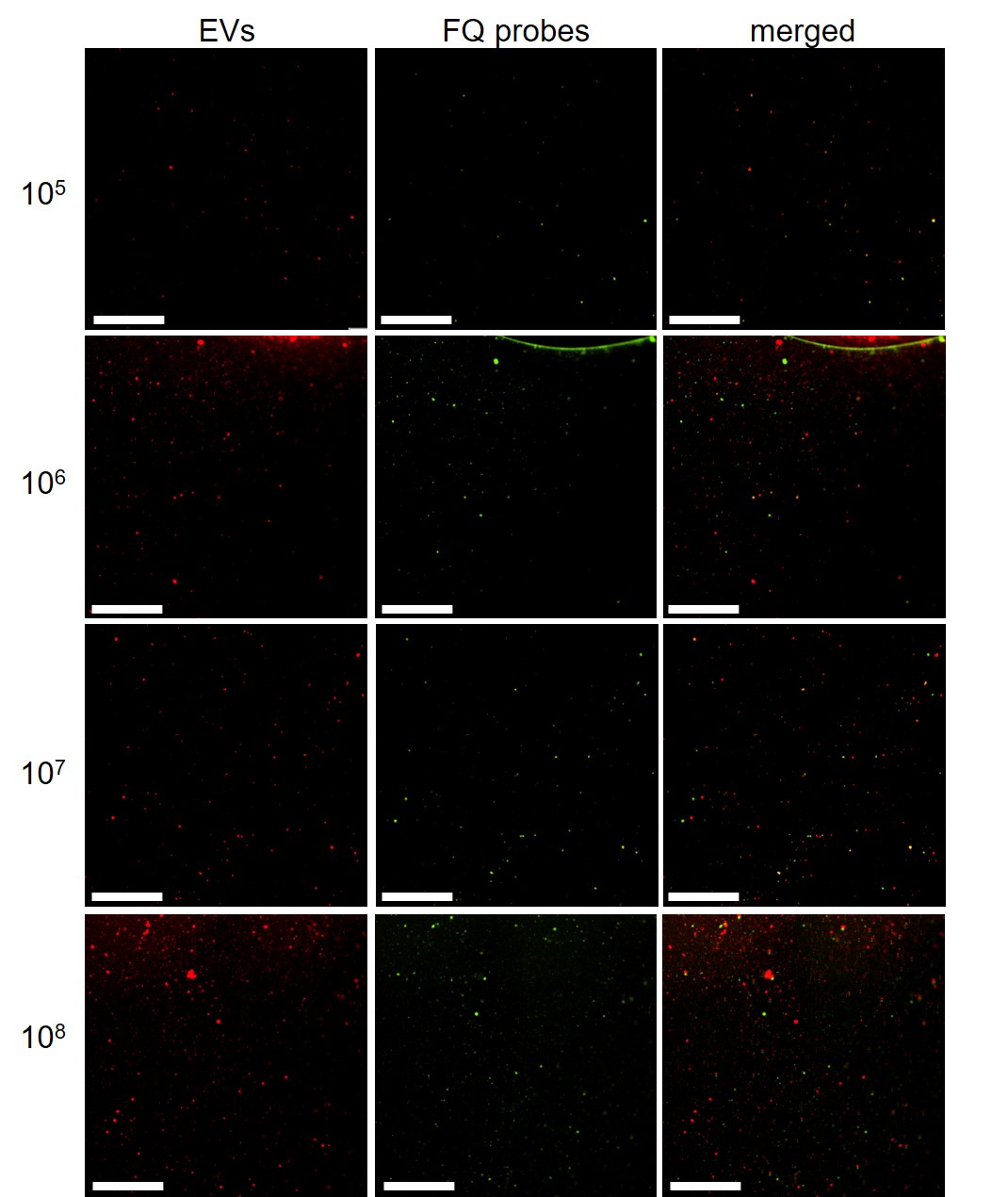

**Figure S2. Fluorescence co-localization analysis.** Representative fluorescence images of PKH26-labeled EVs (red, left), cleaved FQ probes (green, middle), and co-localized signals (right). Scare bar is 100 µm.

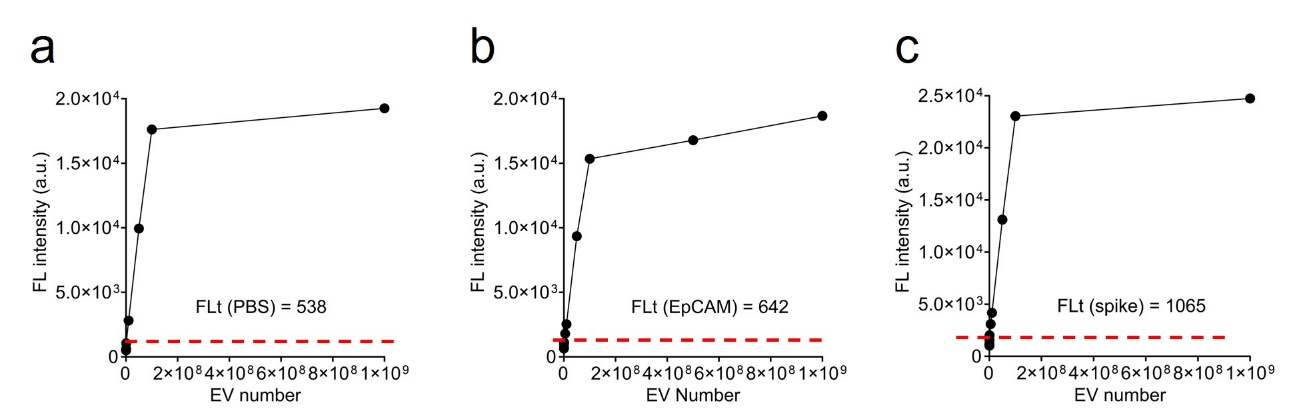

**Figure S3. Fluorescence threshold of mutation detection. a**, fluorescence threshold of L858R detection in H1975 EVs suspended in PBS. **b**, fluorescence threshold of L858R detection in H1975 EVs immunocaptured on surfaces. **c**, fluorescence threshold of L858R detection in H1975 EVs spiked in human plasma.

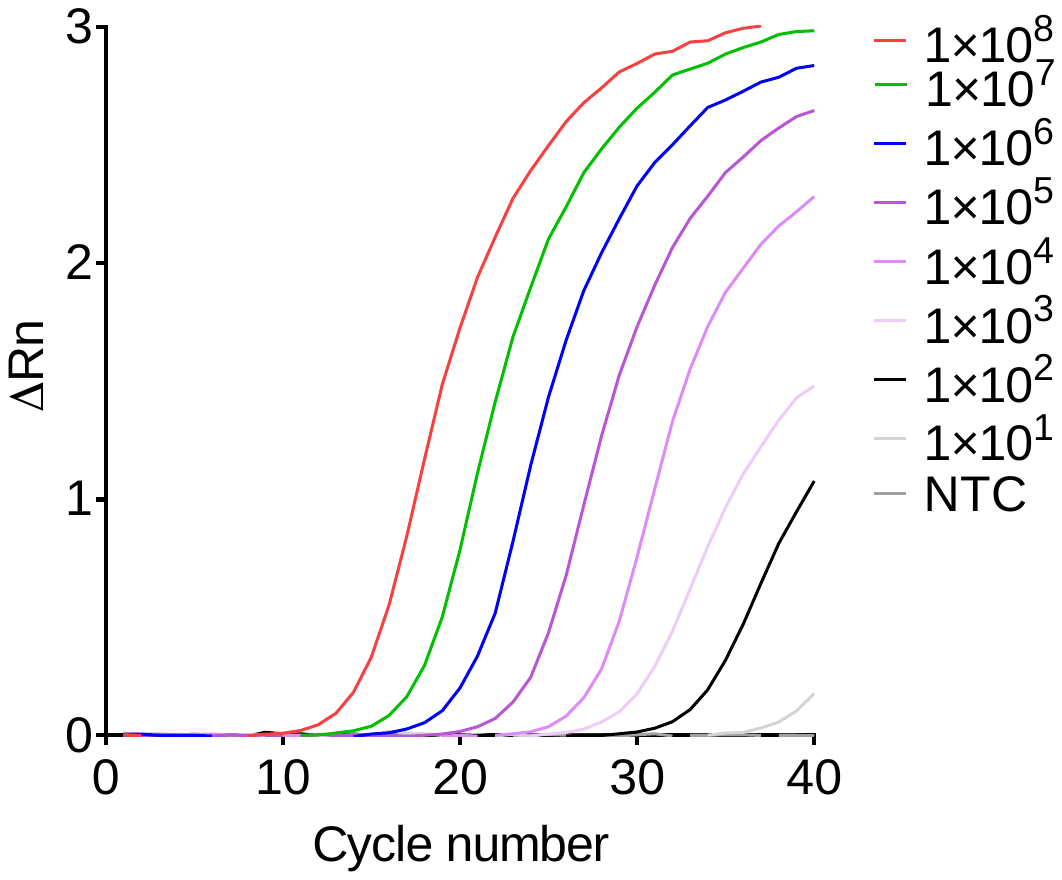

**Figure S4. Detection of L858R by using RT-qPCR**. DNA fragments were extracted from various amount of H1975 EVs followed by RT-qPCR detection.

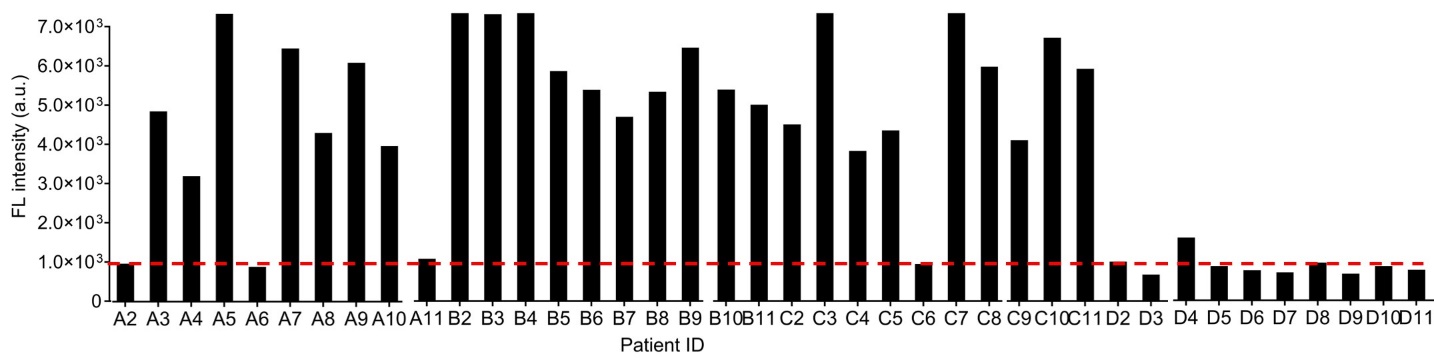

**Figure S5. L858R mutation detection in patient’s plasma EVs.** Thirty patients with stage-IV NSCLC and ten healthy volunteers were enrolled. The red dot line indicates fluorescence threshold of mutation detection.
